# Synergistic olfactory nerve input and cholinergic neuromodulation can activate ERK in rat olfactory bulb vasopressin cells

**DOI:** 10.1101/2025.01.28.635198

**Authors:** Nicolas Reichardt, Lisa Kindler, Esteban Pino, Michael Lukas, Veronica Egger, Hajime Suyama

## Abstract

Social discrimination in rats relies on vasopressin cells (VPCs) intrinsic to the olfactory bulb (OB). We had observed that VPCs responded to electrical stimulation of the olfactory nerve in acute OB slices with inhibitory postsynaptic potentials, and that the neuromodulator acetylcholine could revert these responses to excitation, resulting in action potentials (AP). Moreover, in behaving rats that were exposed to conspecifics, more VPCs were immunopositive to the neural activity marker pERK (pERK^+^ VPC) than in control rats. However, it is unclear whether these two observations, the increased ERK activation *in vivo*, and the generation of APs in the presence of ACh *in vitro*, can be actually mapped onto each other. Here we investigated ERK activation in acute OB slices from transgenic VP-eGFP rats upon either chemical stimulation or tetanic olfactory nerve stimulation. Both KCl and NMDA stimulation resulted in substantial pERK induction across bulbar layers and caused VPC spiking in whole-cell recordings, but only NMDA slightly increased pERK^+^ VPC percentage. Tetanic olfactory nerve stimulation yielded localized, column-like ERK activation of neurons across bulbar layers. The presence of ACh during tetanic stimulation substantially and specifically increased percentages of pERK^+^ VPCs within columns, indicating that columnar pERK^+^ VPCs were preferentially activated by coincident cholinergic neuromodulation and synaptic input.

Our results validate pERK induction as a tool to monitor synaptic VPC excitation in the OB, and imply that depolarization of VPCs alone is insufficient to activate ERK. We propose that synaptically evoked APs are a prerequisite for pERK induction in VPCs.

## Introduction

Detection of neural activity is essential for understanding the function of neural circuits. The relevant methods include immediate observation of activity via electrophysiology, Ca^2+^ imaging and voltage imaging, slightly delayed observation via haemodynamic changes (BOLD signal), and post-hoc observation of the expression of immediate early genes such as c-Fos. The latter technique enables non-invasive studies of neuronal processing in behaving animals, thanks to the time lag between neural activity and expression or induction of markers. However, since these methods report different readouts of neural activity at different detection thresholds, e.g. changes in membrane potential, Ca^2+^ concentration or protein expression, mapping them onto each other is a challenge. Nevertheless, such a mapping is desirable for multi-faceted experimental approaches that combine *in vivo* and in *vitro* assays.

The activity-dependent marker pERK (phosphorylated extracellular signal-related kinase) is part of an intracellular signaling cascade that is involved in e.g. LTP induction in hippocampal CA1 pyramidal neurons *in vitro* (Dudek and Fields 2001). Here, ERK activation is muscimol- (a GABA-A receptor agonist) and CNQX- (an AMPA glutamatergic antagonist) sensitive, and thus dependent on synaptic transmission. Moreover, stimulation below spiking threshold does not activate ERK, supporting the idea that ERK activation is also action potential (AP) dependent. As to sensory areas, both noxious stimuli *in vivo* and electrical activation of C-fiber afferents *in vitro* result in pERK signals in the dorsal horn of the spinal cord (Ji et al. 1999; Kawasaki et al. 2004; Lever et al. 2003) that require the involvement of various glutamate receptors, in particular NMDA receptors (NMDARs; Kawasaki et al. 2004; Xia et al. 1996). In the rat olfactory bulb (OB), odor stimulation (single odorant, urine, conspecific) activates ERK in mitral cells (MC) and granule cells (GC) (Mirich et al. 2004; Suyama et al. 2021). Therefore, pERK is a suitable marker for bulbar neural activation by sensory input. Compared to c-Fos, pERK appears more sensitive, also within the temporal domain, since the peak of its induction happens already within 2 min past the excitatory stimulus (Engholm-Keller et al. 2019; Ji et al. 1999; Xifro et al. 2014).

In this study, we aim to establish a causal link between AP firing and the induction of pERK in the intrinsic vasopressin (VP) system of the rat olfactory bulb. The neuropeptide VP is known to modulate social behavior; in the OB it is acting during social discrimination in rats (reviewed in Suyama et al. 2022). Previously, we showed that the response of bulbar VP cells (VPC) to electrical stimulation of the sensory afferent, the olfactory nerve (ON), in acute OB slices is dominated by inhibitory inputs (Lukas et al., 2019). Furthermore, acetylcholine (ACh) reverts this ON-evoked inhibition to AP firing in the majority of VPCs (Suyama et al. 2021). *In vivo* exposure to different stimuli followed by immunohistochemistry against pERK revealed that social interaction with conspecifics activates more VPCs than control and also activates more cholinergic neurons in the horizontal limb of the diagonal band of Broca than exposure to rat urine (Suyama et al. 2021). Those cholinergic neurons are known to project to the OB (Shute and Lewis 1975; Záborszky et al. 1986), in particular to the glomerular layer, external plexiform layer and MC layer (Kasa et al. 1995). We thus suggested that VPC activation during social interaction is mediated by cholinergic neuromodulation of VPC excitability (Suyama et al. 2021). However, it is not known whether VPC AP firing in OB slices in the presence of ACh and increased ERK activation in VPCs during social interaction are indeed both due to coincident excitation of olfactory inputs **and** ACh release within the OB.

Thus, here we aimed to investigate which type of signal or combinations thereof can indeed activate ERK in bulbar VPCs *in vitro*. To this end, we adopted experimental procedures established previously in the spinal cord (Ji et al. 1999; Kawasaki et al. 2004).

## Materials and methods

### Animals

All experiments were conducted according to national and institutional guidelines for the care and use of laboratory animals, the rules laid down by the EC Council Directive (86/89/ECC), and German animal welfare. Heterozygous VP-eGFP Wistar rats (Ueta et al. 2005) were bred at the University of Regensburg. The light in the rooms was set to an automatic 12 h-cycle (lights on 07:00-19:00).

### Slice preparation

11-19 day-old juvenile rats of either sex were used for the experiments. The rats were deeply anesthetized with isoflurane and quickly decapitated. Horizontal slices (300 µm) were cut in ice-cold carbogenized ACSF (artificial cerebrospinal fluid; in mM: 125 NaCl, 26 NaHCO3, 1.25 NaH2PO4, 20 glucose, 2.5 KCl, 1 MgCl2, and 2 CaCl2) using a vibratome (VT 1200, LEICA, Wetzlar, Germany) and afterward incubated in ACSF at 36 °C for 30 min. The slices were stored at room temperature (∼21 °C) in ACSF for 3 hours before any pERK-inducing assays to reduce unspecific ERK activation due to the slicing procedure (Kawasaki et al. 2004). Prior to electrophysiological recordings slices were stored for at least 1 hour.

### Tetanic olfactory nerve stimulation

Brain slices were placed in a recording chamber as for the electrophysiology. Olfactory nerve (ON) stimulation was performed as described previously (Suyama et al. 2021). Briefly, a stimulation electrode was gently placed in the ON layer of the medial anterior OB. The strength of a single stimulation pulse was 800 µA for 100 µs. Tetanic stimulation at 50 Hz was applied for 6 s, amounting to 300 pulses in total (while three 50 Hz-trains for 1 s spaced at 10 s were used in Kawasaki et al. (2004)).

### Pharmacology and chemical stimulation

The pharmacological agents included acetylcholine (ACh) (100 µM, as in Suyama et al. 2021), KCl (90 mM, as in Kawasaki et al. 2004), and NMDA (100 µM, as in Kawasaki et al. 2004; all Sigma-Aldrich, Darmstadt, Germany). All drugs were diluted at the final concentration in ACSF for bath application during chemical or ON stimulation, or for wash-in during electrophysiological recordings. For the ACh experiments, slices were perfused with ACh starting 10 min before tetanic ON stimulation.

KCl and NMDA stimulations (“chemical stimulation”) were performed by transferring slices into a small chamber with a mesh at the bottom. This chamber was incubated in a beaker filled with ACSF for 5 min, then it was transported into a beaker filled with KCl or NMDA and incubated for 5 min. Following chemical stimulation, the chamber was transferred back to an ACSF beaker and incubated for 10 min (Figure 1A).

**Figure 1.**
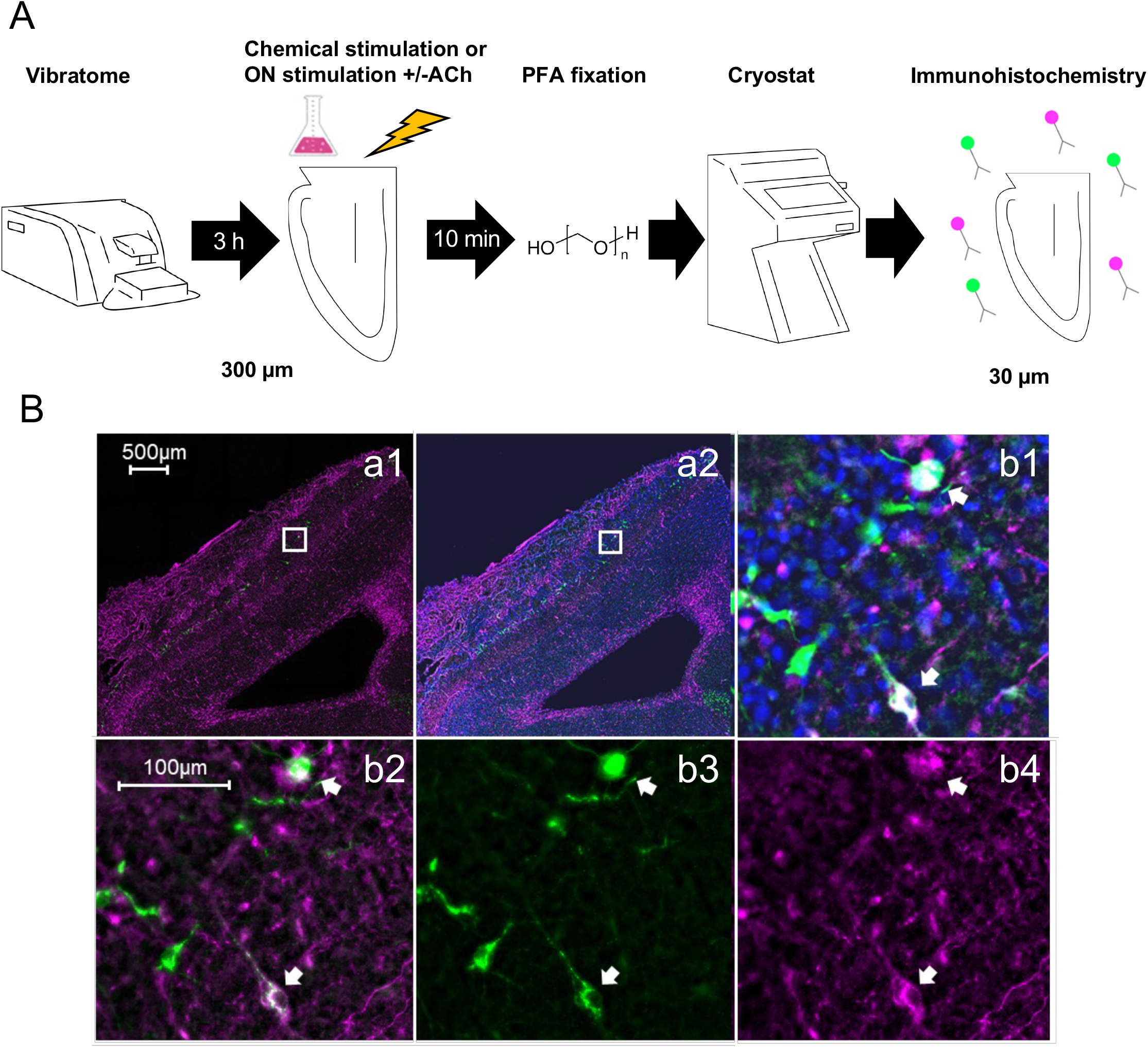
Methods. **A** Experimental design. Acute brain slices of the OB were made with a vibratome, followed by chemical/electrical stimulation and fixation. Slices were further cut into 30-µm thick slices for immunohistochemistry. **B** Representative double and triple labeling of VPCs in control conditions. Green, GFP; Magenta, pERK; Blue: DAPI. a1: overview OB GFP, pERK merge; a2: overview OB GFP, pERK, DAPI merge; Inset: b1: pERK positive VPCs (white arrows) including DAPI merge; b2: pERK positive VPCs merge; b3: GFP channel; b4: pERK channel.

### Cryo-cutting of stimulated slices

10 min after tetanic or chemical stimulation, slices were transferred into a 12-well plate filled with 4 % paraformaldehyde in PBS and fixated at room temperature overnight. Then, PBS was exchanged for 30 % sucrose in PBS to cryoprotect the tissue. To ensure permeation by the immunohistochemical reagents, 300-µm OB slices were cryo-sectioned into 30-µm slices (CM3050 S, LEICA, Wetzlar, Germany) at approx. −20 °C, following Kubota (2021).

### Labeling

Antibody staining against GFP and pERK was performed exactly as previously described (Suyama et al. 2021). In addition, slices were stained for DAPI using DAPI Fluoromount-G (SouthernBiotech, Birmingham, AL, USA).

### Fluorescence microscopy

Fluorescent images of the stained 30-µm slices were obtained as previously described (Suyama et al. 2021) using a DM6 B microscope driven by the software LAS X (LEICA). Z-stack pictures from approx. 6-7 different z-positions per 30-µm slice were used for analysis.

### Cell counting

Immunoreactive VPCs were counted manually across the z-stack pictures of the 30-µm slices using the multi-point tool or cell counter plug-in in Fiji. Double-positive cells were identified by comparing two color channels for pERK (magenta) and GFP (green). DAPI staining (blue) confirmed the presence of a nucleus. The positions of counted cells were recorded. Due to difficulties with cryo-cutting, the number of recovered stained 30-µm slices per original 300-µm slice was variable. For inclusion in the analysis, at least two 30-µm slices per original thick slice containing at least 50 GFP-VPCs had to be recovered. Slices missing a large part of the OB were excluded from the analysis.

In tetanic ON-stimulated slices, immunoreactive cells were counted within all activated regions (denoted “activated”), and outside (denoted “outside”). Activated regions showed a band of pERK^+^ MCs, often with columnar structures of pERK^+^ GCs below. Borders of activated regions were delineated by the positions of the outermost pERK^+^ MCs on either side and drawn perpendicularly to the MCL.

For the experiments following tetanic ON-stimulation or tetanic ON-stimulation combined with ACh wash-in, the numbers of pERK^+^ MCs and pERK^+^ GCs belonging to individual columnar structures were counted as well. Since it was not possible to also count pERK^-^ MCs and GCs, these data could not be normalized. MCs and GCs were identified by their morphological appearance and their localization in the clearly defined MC and GC layers (Halász 1990). MCs were counted manually. For GCs, we used a custom Fiji macro that automatically identified and counted pERK^+^ cells within a certain region of interest (i.e. a column as defined above; see Supplementary methods).

All pictures were analyzed by observers blinded with respect to stimulation groups.

### VPC identification and electrophysiology

VPC identification, whole-cell patch clamp recording and analysis were performed as previously described (Lukas et al. 2019). For identification, a modified Zeiss Axioskop epifluorescence microscope (Carl Zeiss Microscopy, Oberkochen, Germany) with DIC illumination was used. Leaky cells with a holding current above ∼ −10 pA were rejected, as well as recordings that showed a substantial drift in resting V_m_. Pharmacological manipulations were performed via bath application of KCl (90 mM) or NMDA (100 µM), respectively. Traces were analyzed using IGOR Pro 6.37 (Wavemetrics, Portland, USA).

### Statistics

pERK^+^ VPC percentages were computed as the number of pERK^+^ VPCs divided by the number of all VPCs. VPCs and pERK^+^ VPCs were counted in all the 30-µm slices derived from one 300-µm slice.

Statistics were performed with SPSS (ver. 28, IBM, Armonk, NY, USA) and G*Power (ver. 3.1.9.2, Franz Faul, University of Kiel). All statistical analysis performed was non-parametric and two-sided. The used tests are indicated in the text. To compare across groups for chemical stimulation (control, KCl, NMDA) the Kruskal-Wallis test was used, followed by a Dunn-Bonferroni post-hoc test for pairwise comparisons. The Mann-Whitney U test was used for nonpaired and the Wilcoxon Test for paired group comparisons. Effect sizes were calculated using classical Cohen’s d for Mann-Whitney U test, Cohen’s dz for Wilcoxon Tests or Cohen’s f for the Kruskal-Wallis test. Sample sizes were either based on prior studies or on a priori power analysis. Power, effect sizes and sample sizes (for α=0.05) were determined using G*Power (Faul et al. 2007). Details including confidence intervals are given in Table 1. Since the number of recovered 30-µm slices per 300-µm slice differed substantially across experiments (range 2-8), we also weighted the data to check for potential confounding influences. No substantial deviations were observed (see Results). All data are given as mean ± standard deviation.

**Table 1.** Statistical overview.

|  | panel | type of test;<br>all non-parametric | degrees of<br>freedom df | p-<br>value | effect size | power | confidence<br>interval (95%) |
| --- | --- | --- | --- | --- | --- | --- | --- |
| a | 2B | Kruskal-Wallis | 2.0 | 0.010 | f= 0.523 | 0.770 |  |
| b | 2B<sub>1</sub> | Dunn's-Bonferroni | 20.0 | 0.994 | d= 0.344 | 0.120 | [-0.7, 1.4] |
| c | 2B<sub>2</sub> | Dunn's-Bonferroni | 23.0 | 0.158 | d= 0.785 | 0.463 | [-1.9, 0.1] |
| d | 2B<sub>3</sub> | Dunn's-Bonferroni | 23.0 | 0.009 | d= 1.511 | 0.948 | [-1.9, -0.5] |
| e | - | Mann-Whitney-U | 20.9 | 0.001 | d= 1.747 | 0.977 | [-4.2, -1.4] |
| f | 3B<sub>1</sub> | Wilcoxon | 11.4 | 0.013 | dz= 0.875 | 0.804 | [-6.3, -1.4] |
| g | 3B<sub>2</sub> | Mann-Whitney-U | 28.6 | 0.209 | d= 0.494 | 0.255 | [-0.5, 2.1] |
| h | 3B<sub>3</sub> | Mann-Whitney-U | 28.6 | 0.003 | d= 1.257 | 0.909 | [4.1, 12.2] |
| i | 3B<sub>4</sub> | Wilcoxon | 17.1 | <0.001 | dz= 1.743 | 0.999 | [-14.5, -6.2] |
| j | 3C<sub>1</sub> | Mann-Whitney-U | 38.1 | 0.054 | d= 0.522 | 0.347 | [-34.0, 0.1] |
| k | 3C<sub>2</sub> | Mann-Whitney-U | 38.1 | 0.156 | d= 0.483 | 0.305 | [-2.75, 0.5] |

## Results

In order to link our *in vivo* observations using pERK as a neural activity marker to our *in vitro* recordings of ON-evoked VPC spiking in the presence of ACh, we used a cross-over approach in which neural activity *in vitro* was analyzed in terms of ERK activation (Figure 1A). We first established the basal pERK induction level in acute slices (see Methods). The averaged baseline percentage of pERK^+^ VPCs (pERK-positive VPCs) was 1.7 ± 1.3 % (control in Figure 2A, B, N = 11 300-µm slices from 8 rats; weighted mean 1.5 ± 1.2 %; numbers of VPCs per 30-µm slice, see Figure S1).

**Fig 2.**
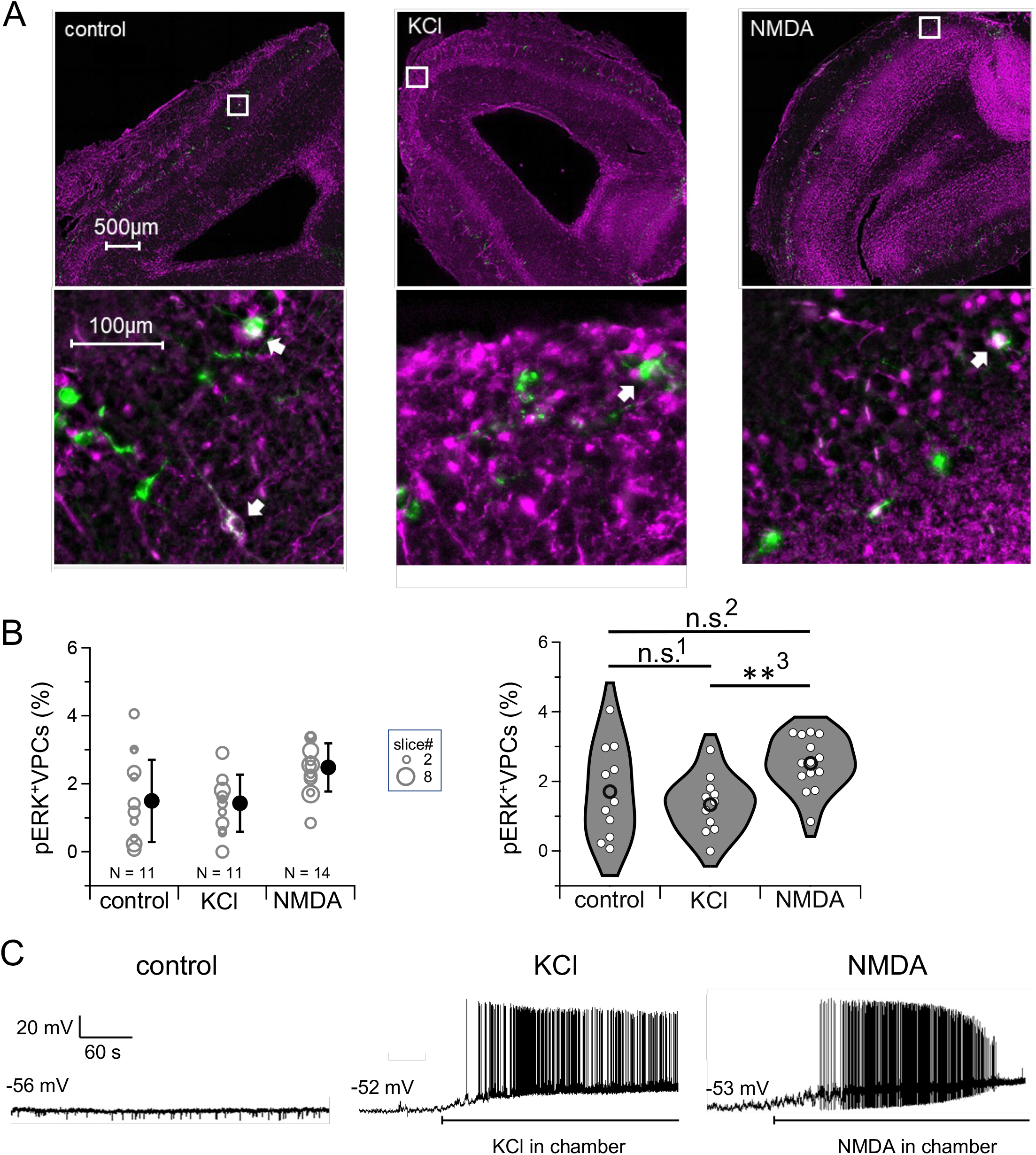
Chemical stimulation – control, KCl, NMDA. **A** Representative examples of pERK labeling in OB slices from control (ACSF), KCl (90 mM), NMDA (100 µM) conditions. Green, GFP; Magenta, pERK. Top: overview. Bottom: inset. **B** Cumulative analysis. Left panel: data points, size scaled with weight, and weighted mean values ± SD. Right panel: non-weighted data points and statistics. **C** Representative whole-cell 6-minute recordings of spontaneous VPC activity for each condition. Initial resting membrane potential indicated at left.

### KCl did not increase ERK activation in VPCs

We first examined whether KCl-induced depolarization can cause VPC spiking and subsequent ERK activation, since Tobin et al. (2010) showed that VP was released from OBs upon stimulation with 50 mM KCl. Moreover, KCl stimulation activated ERK in superficial dorsal horn neurons in spinal cord slices (90mM) (Kawasaki et al. 2004).

In KCl-stimulated OB slices, substantial neuronal pERK signals were found from the GL down to the GCL (Fig. 2A). Intriguingly, however, KCl stimulation did not alter the percentage of pERK^+^ VPCs vs control (KCl: 1.3 ± 0.8 %, N = 11 from 5 rats, p = 0.99 vs control; Kruskal-Wallis with post-hoc Dunn-Bonferroni, see Table 1 and Methods for more information on all statistics; Figure 2A,B, weighted mean 1.4 ± 0.8 %; number of VPCs per 30-µm slice see Supplementary Figure S1A). This finding was surprising because the high concentration of KCl should shift E_K_ to almost 0 mV and thus massively depolarize V_m_. However, there is a very high fraction of inhibitory neurons within the OB, mostly in the GL and GCL (Shepherd 2004), that become activated, as also reflected in the pERK signal. Furthermore, we observed previously that VPCs receive substantial inhibition from glomerular circuits (Lukas et al. 2019). Thus, during KCl stimulation VPCs might receive strong inhibition from juxtaglomerular interneurons.

To test this possibility, we performed whole cell recordings of spontaneous VPC activity in current-clamp and found that upon wash-in of KCl VPCs were depolarized (5 out of 5 cells from 3 animals) and fired APs (3 out of 5 cells, Fig. 2C). Thus depolarization-induced spiking as such is not sufficient to induce pERK in VPCs. Does it then require the activation of synaptic pathways?

### NMDA increases ERK activation in VPCs

In spinal cord dorsal horn neurons, pERK is induced by various neurotransmitters including NMDA (Kawasaki et al. 2004). Moreover, in the hippocampus, an NMDAR antagonist, APV, blocks pERK induction upon electrical stimulation (Dudek and Fields 2001). Therefore, we next examined whether NMDA application can directly induce pERK in bulbar VPCs. In OB slices stimulated with 100 µM NMDA, substantial pERK signals were observed in cell bodies in the GL and the GCL and some dendrites in the EPL (Fig. 2A). NMDA application resulted in a significant increase in the percentage of pERK^+^ VPCs vs the KCl condition (2.5 ± 0.8 %, N = 14 from 8 rats; p = 0.009 Kruskal-Wallis/Dunn-Bonferroni vs KCl, Figure 2A,B; weighted mean 2.5 ± 0.7 %; numbers of VPCs per 30-µm slice Figure S1A), while the comparison with control was not significant, with a tendency for an increase (p = 0.158).

To test the effect of NMDA on VPC electrical activity, we then performed whole-cell recordings from VPCs. Wash-in of NMDA resulted in depolarization (10 of 10) and AP firing (9 of 10) of VPCs (Fig. 2C). We also observed that 4 of the 10 VPCs died within 4 min after the beginning of NMDA application, which might explain the still rather small increase in pERK signal within the VPC population. We conclude that either the NMDAR-dependent pathway of ERK activation is likely to exist in VPCs or that excitatory neurons presynaptic to VPCs such as external tufted cells get substantially activated by NMDA and contribute to synaptically evoked pERK-induction in VPCs (see Discussion).

### Tetanic ON stimulation activates VPCs in the presence of ACh

We next aimed to test whether stimulation of the sensory afferents, the ON, and/or neuromodulation by ACh can increase the percentage of pERK^+^ VPCs. We previously reported that ACh application enables AP firing in response to single ON stimulation in VPCs (Suyama et al. 2021). In line with the pERK study by Kawasaki et al. (2004) that used stimulation of the afferent C-fiber at 50 Hz, we used tetanic ON stimulation (300 times at 50 Hz). Slices were electrically stimulated via the ON either in ACSF (control) or ACh (100 µM). In the ACSF condition, tetanic ON stimulation resulted in regional pERK induction, showing bands of MCs or a columnar pERK signal across all bulbar layers (Figure 3A), reminiscent of earlier reports using retrograde virus tracing (Willhite et al. 2006). These columnar activation patterns most likely reflect the vertical organization of OB cells that belong to the same glomerular unit (e.g. Kikuta et al. 2013); for a quantitative description of these structures see (Jahan et al. BioRxiv 2025).

**Fig 3.**
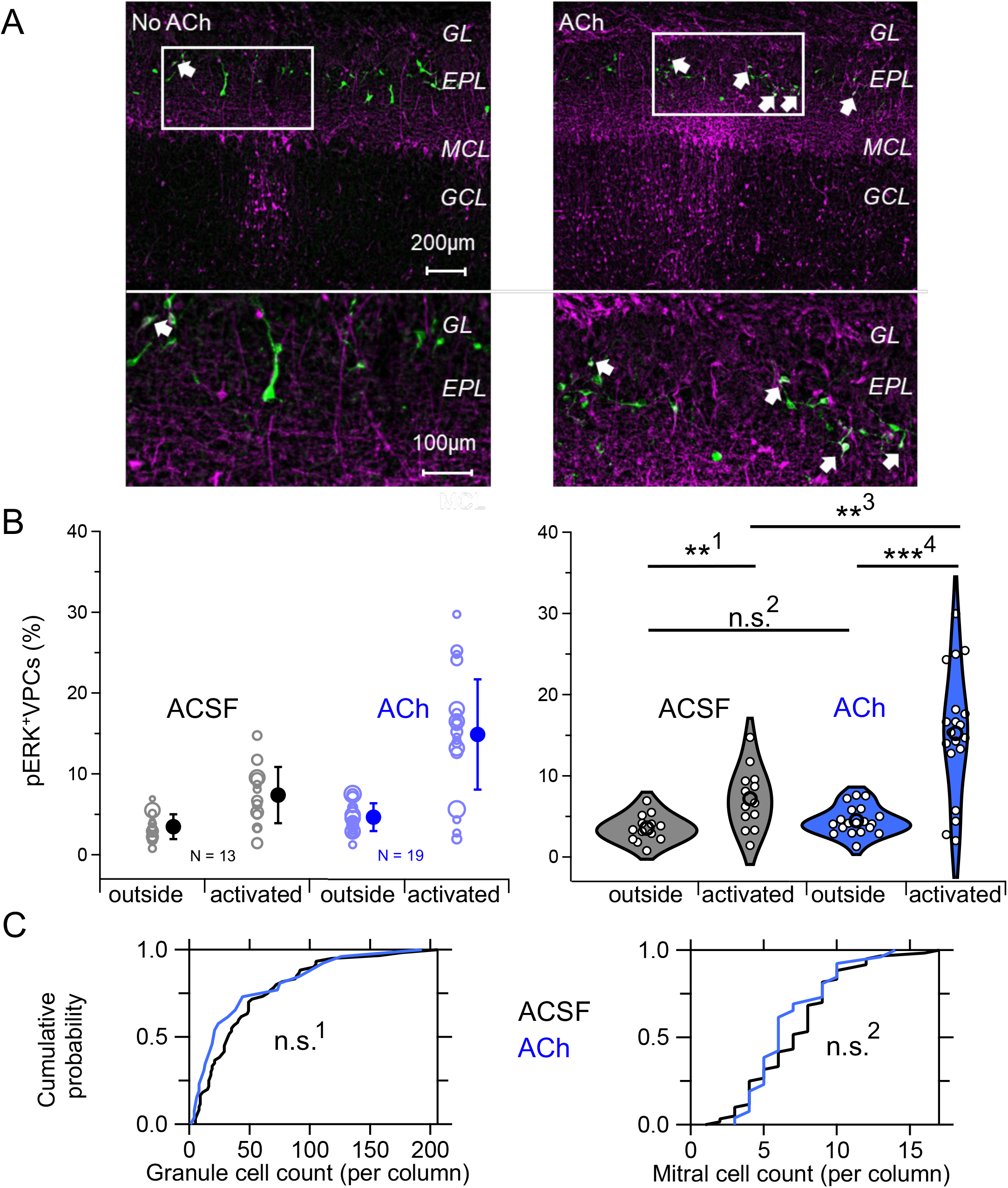
Tetanic olfactory nerve stimulation shows increased VPC-pERK signal in the presence of Ach. **A** Representative examples for control (left) and ACh (right). GL, glomerular layer, EPL, external plexiform layer; MCL, mitral cell layer; GCL, granule cell layer. White arrows indicate pERK^+^ VPCs. Top: overview, bottom: inset. **B** Cumulative analysis of pERK^+^ VPC percentages in and outside of activated regions in the absence and presence of ACh during ON stimulation. Left panel: data points, size scaled with weight as in Figure 2B, and weighted mean values ± SD. Right panel: non-weighted distributions and statistics. **C** Cumulative probability discributions of columnar pERK^+^ cell number (counted in 30-µm slices) for granule cells and mitral cells in the absence (n = 59 columns) and presence (n = 29 columns) of ACh.

Next, we compared the percentage of pERK^+^ VPCs above activated regions and outside of any activated regions (Figure 3A, B). The fractional pERK induction in VPCs within activated regions was significantly higher than outside (Wilcoxon test, p = 0.013, activated: 7.2 ± 3.7 % (weighted 7.4 ± 3.5 %), outside: 3.6 ± 1.7 % (weighted 3.5 ± 1.5%), N = 13 from 8 rats. VPC numbers per 30-µm slice in Figure S1B). The latter pERK^+^ VPC percentage was significantly higher than the basal pERK activity in the control slices without any stimulation (Mann-Whitney test, p = 0.001, see Table 1e). Thus, ON stimulation enhances pERK^+^ induction in VPCs.

In the ACh condition, tetanic ON stimulation also induced pERK in a regional manner (Figure 3A, B). The fractional pERK induction in VPCs within activated regions was significantly and substantially higher than outside (Wilcoxon signed-rank test, p < 0.001, activated/ACh: 15.3 ± 7.7 % (weighted 15.0 ± 6.9 %), outside/ACh: 4.5 ± 1.7 % (weighted 4.8 ± 1.7 %), N = 19 from 10 rats. VPC numbers per 30-µm slice in Figure S1B). Next, we compared the percentages of pERK^+^ VPCs within activated regions in the presence and absence of ACh. For the activated/ACh condition, the mean percentage of pERK^+^ VPCs was significantly higher than that for the activated/ACSF condition (Mann-Whitney test, p = 0.003). There was no increase in pERK^+^ VPCs for the outside/ACh condition compared to outside/ACSF (Mann-Whitney test, p = 0.209).

Therefore, we conclude that the coincident activation of ON input and the neuromodulation by ACh can trigger pERK induction in VPCs more efficiently than ON input alone.

### Enhancing effect of ACh is specific to VPCs

ACh receptors are known to be expressed across various OB cell populations (Le Jeune et al. 1995). We asked whether ACh might also modulate pERK induction in other bulbar neuron types, since in both MCs and GCs firing rates are subject to modulation via cholinergic pathways (Castillo et al. 1999). Since pERK^+^ GCs mostly occurred within columnar structures, we counted pERK^+^ MCs and GCs within columns (see Methods). The numbers of pERK^+^ MCs and GCs within activated columns were not significantly different between the ACh condition and the ACSF condition (Figure 3C; Mann-Whitney U test; for the numbers of pERK^+^ GCs p = 0.054, column/ACSF: 48.8 ± 36.7, n= 59 columns in 26 slices from 16 rats; column/ACh 27.3 ± 38.5, n= 29 columns in 16 slices from 9 rats; for the numbers of pERK^+^ MCs, p = 0.156, column/ACSF: 7.3 ± 2.8, n= 59 in 26 slices from 16 rats; column/ACh : 6.1 ± 2.0, n= 29 in 16 slices from 9 rats).

We conclude that the observed effect of ACh on pERK induction in VPCs is unlikely to be due to a general disinhibition of the OB by ACh, but rather either a VPC-specific effect or possibly an effect on the glomerular neurons presynaptic to VPCs.

## Discussion

Our experiments were initially intended to validate pERK as a sensitive marker of spiking activity in OB neurons, in particular in VPCs, since we previously observed that VPCs spike only upon coincident sensory input and ACh neuromodulation (Suyama et al. 2021).

However, we observed no difference in the percentage of pERK^+^ VPCs between the KCl condition and the ACSF condition, even though VPCs were firing in the KCl condition (Figure 2). In the spinal cord, KCl stimulation was shown to activate ERK predominantly in superficial dorsal horn neurons but not in all of them (Kawasaki et al. 2004). Therefore, direct depolarization of neurons does not necessarily activate ERK, and this seems to also be the case for VPCs. Another possibility is that all VPCs died upon KCl treatment and therefore no pERK induction was observed. This would indicate a particular vulnerability of VPCs since KCl treatment otherwise resulted in broad pERK expression in the OB slices.

The NMDAR pathway is known to play an important role in ERK activation in different types of neurons (Dudek and Fields 2001; Kawasaki et al. 2004; Xia et al. 1996), most likely via pathways different from the ones downstream of KCl stimulation (e.g. Xifro et al. 2014). Here, NMDA activated ERK in VPCs significantly more than the KCl condition in the OB; the observed increase by NMDA versus control was not significant (Figure 2). ON stimulation evokes IPSPs in VPCs *in vitro*. However, upon blockade of inhibition, ON stimulation evokes EPSPs. Thus, VPCs receive excitatory inputs that are masked by the dominant inhibitory inputs (Lukas et al. 2019). Our results could be explained either by the presence of NMDARs on VPCs and thus a direct excitation or by a strong NMDAR component in the cells providing the excitatory inputs to VPCs such as external tufted cells – or both.

Similar to KCl stimulation effects, a subset of VPCs died upon NMDA stimulation and therefore pERK induction in these cells could have been obliterated. Again, this observation indicates a particular vulnerability of VPCs for excitotoxicity, since NMDA treatment otherwise resulted in broad pERK induction in our OB slices - e.g. in the GCL, in line with the major role of NMDARs in GC excitation (Isaacson and Strowbridge 1998; Mueller and Egger 2020; Najac et al. 2015). These chemical stimulation data indicate that depolarization of VPCs alone is not sufficient to induce pERK, whereas excitatory synaptic transmission involving NMDARs is more efficient.

Tetanic ON-stimulation resulted in regional ERK activation of neurons, often in a columnar manner (see Jahan et al. BioRxiv 2025 for details). Due to the tortuous apical VPC dendrite (Lukas et al. 2019), an unambiguous association between a given VPC and an activated region is not always possible and therefore some pERK^+^ VPCs might have been wrongly assigned. ON stimulation alone did already slightly increase the percentage of pERK^+^ VPCs in general (outside vs control), perhaps due to activation of long-range excitatory glomerular networks (e.g. Aungst et al. 2003), which in turn might provide synaptic excitation to some VPCs. Such excitation would be enhanced for VPCs within an activated region.

Most importantly, in the presence of ACh ON stimulation substantially increased the percentage of pERK^+^ VPCs compared to ON stimulation alone. Since ACh enables VPCs to spike in response to single ON stimulation *in vitro* (Suyama et al., 2021) and since depolarization alone does not trigger pERK induction (Figure 2), we hereby have demonstrated a correlation between synaptically evoked AP firing and pERK signaling in VPCs. Notably, pERK is induced within a limited subset of AP-firing VPCs, since in the previous study 65% of VPCs were shown to fire APs upon ON stimulation in the presence of ACh, whereas the mean percentage of pERK^+^ VPCs observed here is just 15%. Thus, pERK is unlikely to report single VPC APs.

ACh was observed to enhance the activity of excitatory neurons in the OB (Elaagouby et al. 1991; Rothermel et al. 2014). However, in our hands ACh did not increase the intrinsic excitability of VPCs (Suyama et al. 2021) and in the present study we observed a slightly, but not significantly increased percentage of pERK^+^ VPCs between outside/ACSF and the outside/ACh condition (Figure 3B). Therefore, ACh is unlikely to directly activate the ERK pathway in VPCs. We suggest, then, that the observed ACh effects on VPCs upon tetanic ON stimulation are due to cholinergic modulations of the glomerular network that evoke synaptic AP firing and therewith trigger the ERK signaling cascade in VPCs.

In our previous study (Suyama et al. 2021), social interaction activated ERK in ∼50 % of bulbar VPC, while only ∼15 % of VPCs were pERK^+^ in the activated/ACh condition investigated here. This difference in pERK^+^ VPC percentage could be simply due the different stimulation of sensory afferents, i.e. by complex body odors (Singer et al. 1997) likely to cause broad activation, versus spatially limited electrical stimulation of single or a few ON fascicles. Moreover, in the *in-vivo* situation other centrifugal neuromodulators such as noradrenaline (McLean et al. 1989) and serotonin (McLean and Shipley 1987) might also be released in the OB during social interaction in addition to ACh, possibly enhancing VPC activation as we have demonstrated before *in vitro* (Suyama et al. 2021). Therefore, those centrifugal neuromodulators might act synergistically on VPCs *in vivo*.

In conclusion, the previously missing link between ERK activation during social interaction and evoked APs in the ACh condition in VPCs *in vitro* is demonstrated here. Our results imply that cholinergic signaling plays a permissive role in activating ERK in VPCs during social interaction by strengthening VPC synaptic excitation.

## Acknowledgements

Funding: The work was supported by the German research foundation, DFG (LU2164/1-1, GRK 2174, FOR5424: LU2164/2-1, EG 135/12-1). We thank Anne Pietryga-Krieger and Ellen Fröhlich for technical assistance.

## Author contributions

NR performed experiments, analyzed data, prepared figures, edited and revised manuscript.

LK performed experiments, analyzed data.

EP performed experiments, analyzed data.

ML conceived and designed research, performed experiments.

VE conceived and designed research, analyzed data, prepared figures, edited and revised manuscript.

HS conceived and designed research, analyzed data, drafted manuscript, prepared figures, edited and revised manuscript.

All authors approved the final version of the manuscript.

---

**Supplementary Figure 1.**
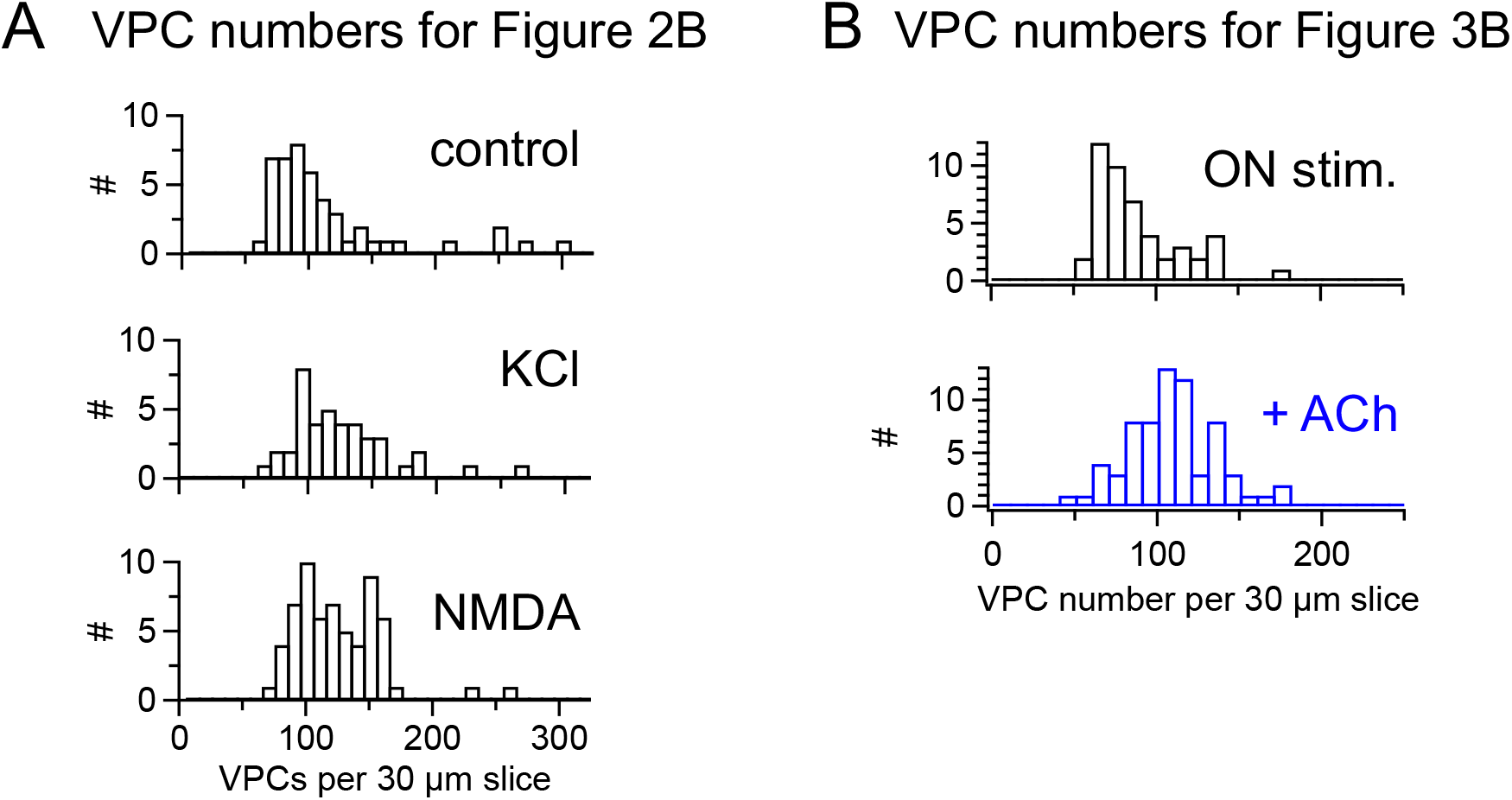
Vasopressin cell numbers across conditions. Counts of VPCs in the respective conditions across 30-µm slices. **A** Control: 114 ± 55, n = 47 slices, KCl: 124 ± 41, n = 41 slices, NMDA: 124 ± 34, n = 62 slices **B** ON stimulation: 87 ± 26, n = 47 slices; ON stimulation + ACh: 107 ± 27, n = 68 slices.

## Supplementary Methods

### Fiji Macro for granule cell counting

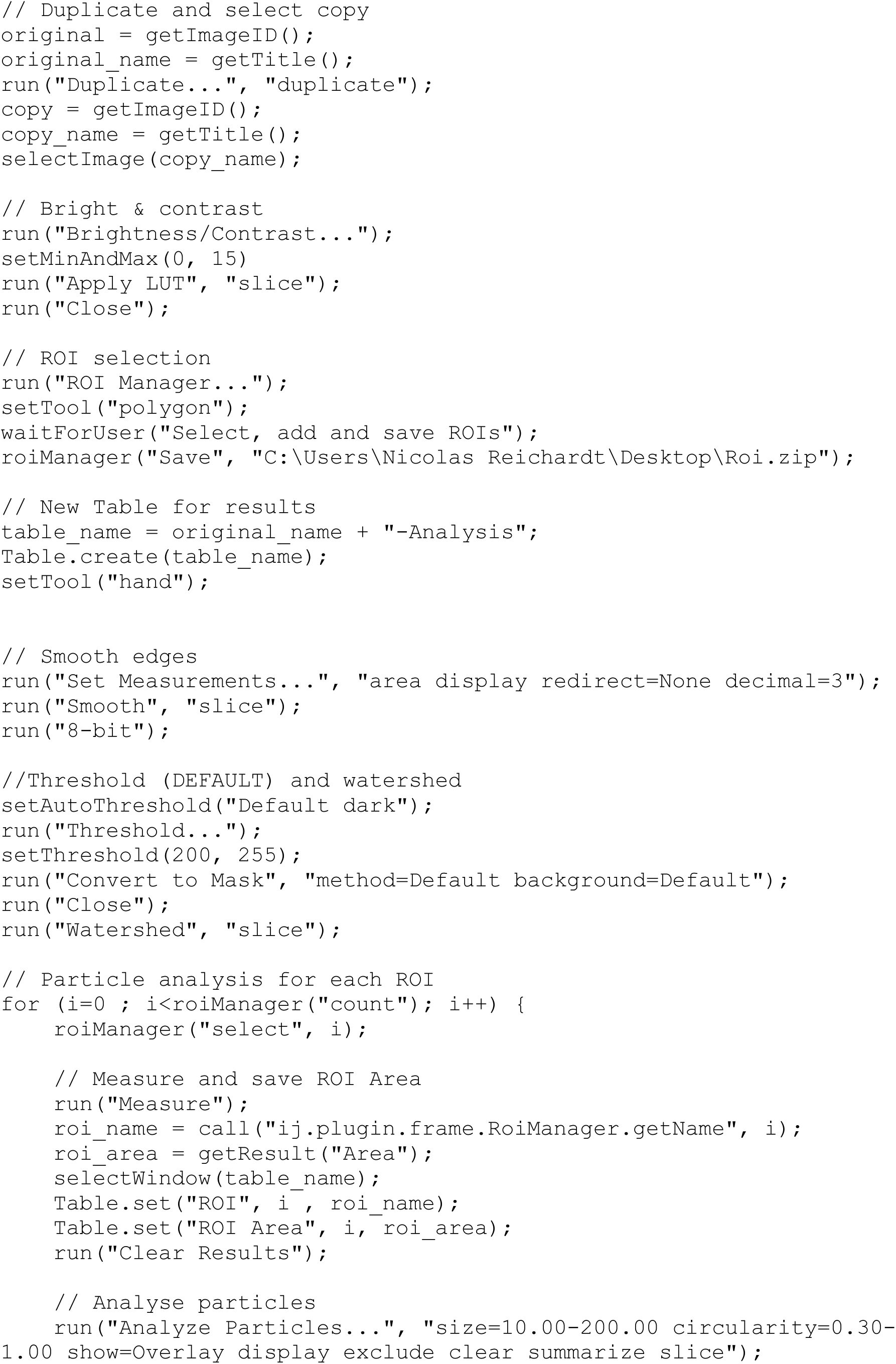
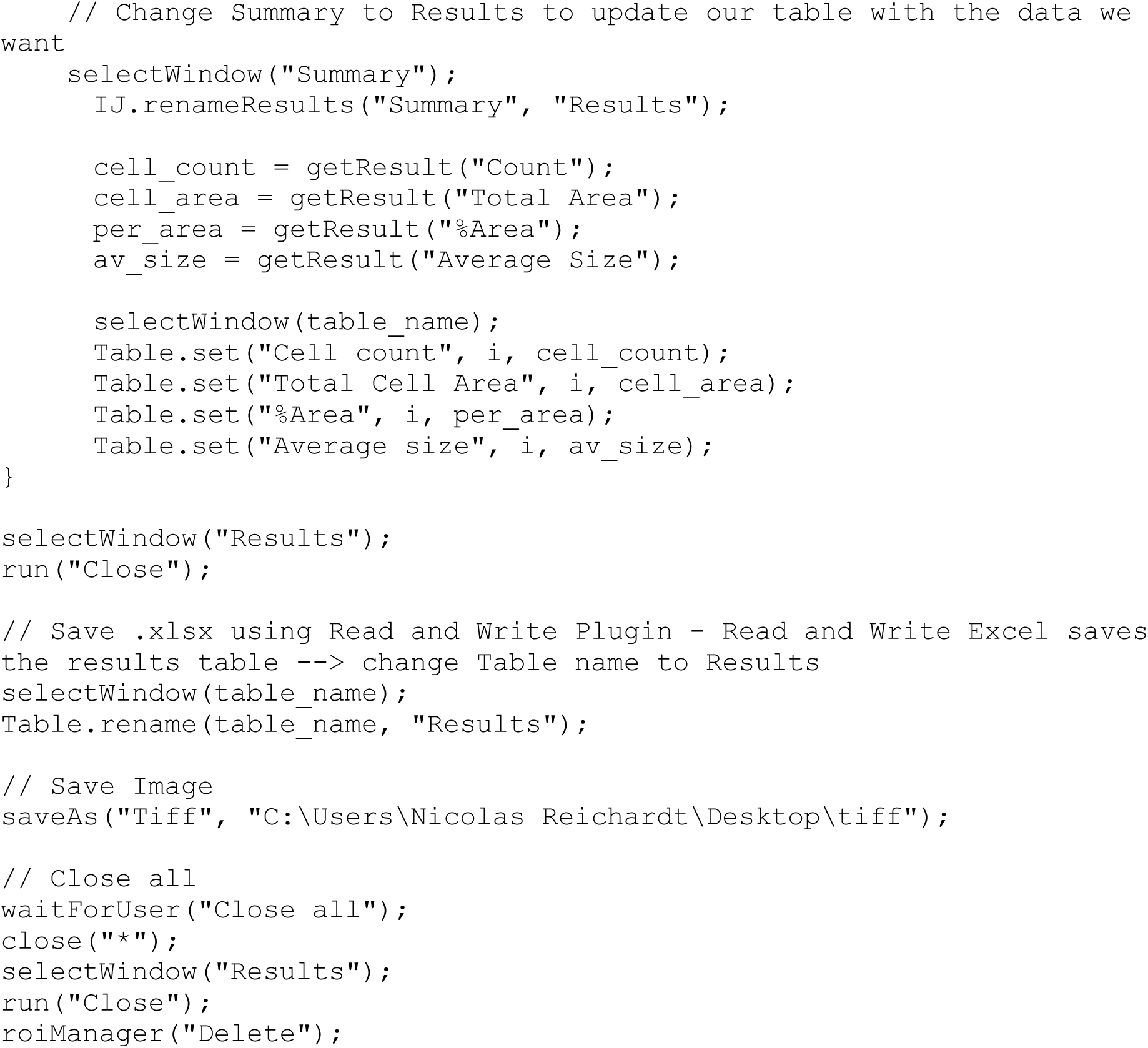

## References

Aungst JL, Heyward PM, Puche AC, Karnup SV, Hayar A, Szabo G, Shipley MT. Centre– surround inhibition among olfactory bulb glomeruli. Nature 426: 623–629, 2003.

Dudek SM, Fields RD. Mitogen-Activated Protein Kinase/Extracellular Signal-Regulated Kinase Activation in Somatodendritic Compartments: Roles of Action Potentials, Frequency, and Mode of Calcium Entry. J Neurosci 21: RC122–RC122, 2001.

Elaagouby A, Ravel N, Gervais R. Cholinergic modulation of excitability in the rat olfactory bulb: Effect of local application of cholinergic agents on evoked field potentials. Neuroscience 45: 653–662, 1991.

Engholm-Keller K, Waardenberg AJ, Müller JA, Wark JR, Fernando RN, Arthur JW, Robinson PJ, Dietrich D, Schoch S, Graham ME. The temporal profile of activity-dependent presynaptic phospho-signalling reveals long-lasting patterns of poststimulus regulation. PLOS Biol 17: e3000170, 2019.

Faul F, Erdfelder E, Lang A-G, Buchner A. G*Power 3: A flexible statistical power analysis program for the social, behavioral, and biomedical sciences. Behav Res Methods 39: 175–191, 2007.

Halász N. The vertebrate olfactory system : chemical neuroanatomy, function and development [Online]. Akadémiai Kiadó.https://cir.nii.ac.jp/crid/1130000793912047360 [22 Nov. 2024].

Isaacson JS, Strowbridge BW. Olfactory Reciprocal Synapses: Dendritic Signaling in the CNS. Neuron 20: 749–761, 1998.

Jahan I, Kindler L, Reichardt N, Suyama H, Egger V. Local olfactory nerve stimulation triggers patchy activation of columns across rat olfactory bulb. BioRxiv Jan 2025

Ji R-R, Baba H, Brenner GJ, Woolf CJ. Nociceptive-specific activation of ERK in spinal neurons contributes to pain hypersensitivity. Nat Neurosci 2: 1114–1119, 1999.

Kasa P, Hlavati I, Dobo E, Wolff A, Joo F, Wolff JR. Synaptic and non-synaptic cholinergic innervation of the various types of neurons in the main olfactory bulb of adult rat: Immunocytochemistry of choline acetyltransferase. Neuroscience 67: 667–677, 1995.

Kawasaki Y, Kohno T, Zhuang Z-Y, Brenner GJ, Wang H, Meer CVD, Befort K, Woolf CJ, Ji R-R. Ionotropic and Metabotropic Receptors, Protein Kinase A, Protein Kinase C, and Src Contribute to C-Fiber-Induced ERK Activation and cAMP Response Element-Binding Protein Phosphorylation in Dorsal Horn Neurons, Leading to Central Sensitization. J Neurosci 24: 8310–8321, 2004.

Kikuta S, Fletcher ML, Homma R, Yamasoba T, Nagayama S. Odorant Response Properties of Individual Neurons in an Olfactory Glomerular Module. Neuron 77: 1122–1135, 2013.

Le Jeune H, Aubert I, Jourdan F, Quirion R. Comparative laminar distribution of various autoradiographic cholinergic markers in adult rat main olfactory bulb. J Chem Neuroanat 9: 99–112, 1995.

Lever IJ, Pezet S, McMahon SB, Malcangio M. The signaling components of sensory fiber transmission involved in the activation of ERK MAP kinase in the mouse dorsal horn. Mol Cell Neurosci 24: 259–270, 2003.

Lukas M, Suyama H, Egger V. Vasopressin Cells in the Rodent Olfactory Bulb Resemble Non-Bursting Superficial Tufted Cells and Are Primarily Inhibited upon Olfactory Nerve Stimulation. eNeuro 6, 2019.

McLean JH, Shipley MT. Serotonergic afferents to the rat olfactory bulb: I. Origins and laminar specificity of serotonergic inputs in the adult rat. J Neurosci 7: 3016–3028, 1987.

McLean JH, Shipley MT, Nickell WT, Aston-Jones G, Reyher CKH. Chemoanatomical organization of the noradrenergic input from locus coeruleus to the olfactory bulb of the adult rat. J Comp Neurol 285: 339–349, 1989.

Mirich JM, Illig KR, Brunjes PC. Experience-dependent activation of extracellular signal-related kinase (ERK) in the olfactory bulb. J Comp Neurol 479: 234–241, 2004.

Mueller M, Egger V. Dendritic integration in olfactory bulb granule cells upon simultaneous multispine activation: Low thresholds for nonlocal spiking activity. PLOS Biol 18: e3000873, 2020.

Najac M, Diez AS, Kumar A, Benito N, Charpak S, Jan DDS. Intraglomerular Lateral Inhibition Promotes Spike Timing Variability in Principal Neurons of the Olfactory Bulb. J Neurosci 35: 4319–4331, 2015.

Rothermel M, Carey RM, Puche A, Shipley MT, Wachowiak M. Cholinergic Inputs from Basal Forebrain Add an Excitatory Bias to Odor Coding in the Olfactory Bulb. J Neurosci 34: 4654–4664, 2014.

Shepherd GM. The Olfactory Bulb. In: Sensory Systems: II: Senses Other than Vision, edited by Wolfe JM. Birkhäuser, p. 66–68.

Shute CCD, Lewis PR. Cholinergic pathways. Pharmacol Ther [B] 1: 79–87, 1975.

Singer AG, Beauchamp GK, Yamazaki K. Volatile signals of the major histocompatibility complex in male mouse urine. Proc Natl Acad Sci 94: 2210–2214, 1997.

Suyama H, Egger V, Lukas M. Top-down acetylcholine signaling via olfactory bulb vasopressin cells contributes to social discrimination in rats. Commun Biol 4: 1–17, 2021.

Suyama H, Egger V, Lukas M. Mammalian social memory relies on neuromodulation in the olfactory bulb. Neuroforum 28: 143–150, 2022.

Tobin VA, Hashimoto H, Wacker DW, Takayanagi Y, Langnaese K, Caquineau C, Noack J, Landgraf R, Onaka T, Leng G, Meddle SL, Engelmann M, Ludwig M. An intrinsic vasopressin system in the olfactory bulb is involved in social recognition. Nature 464: 413–417, 2010.

Ueta Y, Fujihara H, Serino R, Dayanithi G, Ozawa H, Matsuda K, Kawata M, Yamada J, Ueno S, Fukuda A, Murphy D. Transgenic Expression of Enhanced Green Fluorescent Protein Enables Direct Visualization for Physiological Studies of Vasopressin Neurons and Isolated Nerve Terminals of the Rat. Endocrinology 146: 406–413, 2005.

Willhite DC, Nguyen KT, Masurkar AV, Greer CA, Shepherd GM, Chen WR. Viral tracing identifies distributed columnar organization in the olfactory bulb. Proc Natl Acad Sci 103: 12592–12597, 2006.

Xia Z, Dudek H, Miranti CK, Greenberg ME. Calcium Influx via the NMDA Receptor Induces Immediate Early Gene Transcription by a MAP Kinase/ERK-Dependent Mechanism. J Neurosci 16: 5425–5436, 1996.

Xifro Collsamata FX, Miñano Molina AJ, Saura Antolín C, Rodríguez Álvarez J. Ras activation is a key event in activity-dependent survival of cerebellar granule neurons. J Biol Chem 289: 8462–847, 2014.

Záborszky L, Carlsen J, Brashear HR, Heimer L. Cholinergic and GABAergic afferents to the olfactory bulb in the rat with special emphasis on the projection neurons in the nucleus of the horizontal limb of the diagonal band. J Comp Neurol 243: 488–509, 1986.

